# Switch-1 Instability at the Active Site Decouples ATP Hydrolysis from Force Generation in Myosin II

**DOI:** 10.1101/2020.09.10.292169

**Authors:** Benjamin C. Walker, Claire E. Walczak, Jared C. Cochran

## Abstract

Myosin active site elements (i.e. switch-1) bind both ATP and a divalent metal to coordinate ATP hydrolysis. ATP hydrolysis at the active site is linked via allosteric communication to the actin polymer binding site and lever arm movement, thus coupling the free energy of ATP hydrolysis to force generation. How active site motifs are functionally linked to actin binding and the power stroke is still poorly understood. We hypothesize that destabilizing switch-1 movement at the active site will negatively affect the tight coupling of ATP hydrolysis to force production. Using a metal-switch system, we tested the effect of interfering with switch-1 coordination of the divalent metal cofactor on force generation. We found that while ATPase activity increased, motility was inhibited. Our results demonstrate that a single atom change that affects the switch-1 interaction with the divalent metal directly regulates actin binding and force generation. Even slight modification of the switch-1 divalent metal coordination can decouple ATP hydrolysis from motility. Switch-1 movement is therefore critical for both structural communication with the actin binding site, as well as coupling the energy of ATP hydrolysis to force generation.

## INTRODUCTION

Myosins and kinesins are structurally related cytoskeletal molecular motors that use the free energy of ATP hydrolysis to produce force along filament tracks (filamentous actin or microtubules) and play essential roles for a diverse range of cellular processes. The conversion of chemical energy at their active sites to mechanical energy at the filament binding interface and force producing elements is facilitated by conformational changes in their structurally similar central core (Kull & Endow, 2002; Kull, Sablin, Lau, Fletterick, & Vale, 1996; Kull, Vale, & Fletterick, 1998; Sack, Kull, & Mandelkow, 1999; Vale & Milligan, 2000).

Myosins and kinesins have three primary active site elements in their central core that are necessary for nucleotide and divalent metal binding (usually Mg^2+^) to coordinate efficient ATP hydrolysis (Vale, 1996). The P-loop (Walker A motif) binds nucleotide and coordinates Mg^2+^, and the switch-1 and switch-2 (Walker B motif) motifs sense and respond to the absence/presence of the γ-phosphate of the bound nucleotide via direct interaction with the nucleotide and Mg^2+^ (Vale & Milligan, 2000). Mg^2+^ at the active site is coordinated by the hydroxyl group of the serine/threonine residue in the P-loop, the oxygen from the β-phosphate of the bound nucleotide, and two water molecules. The remaining two ligand elements necessary to satisfy the octahedral coordination geometry of Mg^2+^ depend on the specific hydrolase and nucleotide state (nucleoside diphosphate or nucleoside triphosphate). Therefore, the metal ion anchors a network of important interactions in the enzyme that are necessary for efficient nucleotide hydrolysis.

For myosins and kinesins to produce work, movement of switch-1 and switch-2 loops at the active site are linked to the filament binding interface and force generating elements via structural changes in the central core (Vale & Milligan, 2000). These conformational changes modulate motor-filament affinity and movement of force producing elements, such as the lever arm in myosin and the neck linker in kinesin, that allow for motor motility. Although kinesin and myosins share significant structural similarities, their thermodynamic and kinetic mechanisms differ. In myosins, ATP binding leads to rapid actomyosin dissociation followed by ATP hydrolysis (Kurzawa & Geeves, 1996); while in kinesins, ATP binding does not dissociate the tight kinesin-microtubule complex, and hydrolysis occurs before detachment (Gilbert, Webb, Brune, & Johnson, 1995). However, in both enzymes the rate limiting steps are stimulated by filament binding (Cochran, 2015).

Force generation by actomyosin occurs via cyclic interactions of filamentous actin (F-actin) and myosin that are mechanochemically coupled to its ATP hydrolysis cycle (Lymn & Taylor, 1971). How active site motifs are mechanochemically coupled to actin binding and the power stroke is an area of active research. ATP binding to the active site results in closure of the nucleotide pocket through movements of the switch-1, switch-2, and P-loop motifs (Bobkov, Sutoh, & Reisler, 1997; Hiratsuka, 1994). Closure of the switch-1 loop onto the γ-phosphate of ATP is directly coupled to opening of the actin binding cleft and weakening of filament affinity (Conibear, Bagshaw, Fajer, Kovacs, & Malnasi-Csizmadia, 2003; Holmes, Angert, Kull, Jahn, & Schroder, 2003; Naber, Malnasi-Csizmadia, Purcell, Cooke, & Pate, 2010). Switch-2 closure, on the other hand, is coupled to the recovery stroke of the lever arm when dissociated from F-actin (Fischer, Windshugel, Horak, Holmes, & Smith, 2005; Koppole, Smith, & Fischer, 2007). Closure of the switches around ATP leads to rapid hydrolysis and subsequent rebinding of F-actin. How switch loop opening and product release of actomyosin are coupled to force generation is controversial and poorly understood (Gyimesi et al., 2008; Llinas et al., 2015; Muretta, Rohde, Johnsrud, Cornea, & Thomas, 2015; Woody, Winkelmann, Capitanio, Ostap, & Goldman, 2019).

In a previous study, we were able to control the enzymatic activity and F-actin binding of a non-motile truncation of *Dictyostelium discoideum* myosin II (M761) by substituting the divalent metal coordinating switch-1 serine (S237) with cysteine, which diminishes its interaction with the Mg^+2^ metal, and thus inhibits its ATPase activity (Cochran, Thompson, & Kull, 2013). Substituting Mg^+2^ with Mn^+2^, which strongly interacts with cysteine, restores metal interaction, ATPase activity, and actin binding. Hence exchanging divalent metals provided a direct and experimentally reversible link between switch-1 and the actin binding cleft. Interestingly, the basal ATPase rate of wild type (WT) myosin in the presence of divalent metals other than Mg^2+^ is very high, which is similar to the high basal rates observed with the S237C mutant (Cochran et al., 2013). In addition, substituting Ca^2+^ leads to high basal ATPase rates that were inhibited in the presence of F-actin for both WT and S237C. This suggests that perturbing the switch-1 and divalent metal coordination leads to substantial structural instability at the active site with switch loops rapidly opening and closing for ATP hydrolysis and product release. How interference with switch-1 at the active site affects force generation is unknown.

Given that switch loop movement and product release are tightly coupled to the power stroke and force generation, we hypothesized that destabilizing switch-1 movement would negatively affect the tight coupling of ATP hydrolysis to force production that results in motility. To test this hypothesis, we fused two α-actinin repeats (2R) that function as an artificial lever arm (Anson, Geeves, Kurzawa, & Manstein, 1996) to M761 and M761(S237C), the truncated myosin II constructs used in our previous study (Cochran et al., 2013). Cosedimentation and steady-state ATPase assays confirmed that the α-actinin repeats did not change the behavior of the motor constructs. F-actin gliding assays indicated that wild-type (WT) myosin motility was reduced in the presence of Mn^2+^ compared to Mg^2+^, and absent in the presence of Ca^2+^. On the other hand, the mutant failed to produce motility under all divalent metal conditions despite robust ATPase activity in the gliding chambers. Together our results support a model whereby switch-1 coordination of the divalent metal at the active site directly regulates actin binding and force generation. Even slight modification of the switch-1 and divalent metal interaction has significant effects on ATP hydrolysis and can decouple hydrolysis from force generation. Switch-1 movement is therefore critical for allosteric communication with the F-actin binding cleft, as well as for coupling ATP hydrolysis to force generation.

## RESULTS AND DISCUSSION

### The M761-2R S237C mutant can rescue F-actin release in the presence of MnATP

To study the effects of divalent metal coordination of nucleotide binding on myosin force production, we mutated serine 237 to cysteine (S237C) in myosin M761 (S1dC fragment of *Dictyostelium discoideum* myosin II) containing two α-actinin repeats (M761-2R), which act as lever arms to allow for motility. The WT and S237C constructs were expressed with a C-terminal 8x HIS purification tag in *D. discoideum*. Purification of the WT and S237C constructs yielded approximately 0.3 mg and 0.1 mg per liter of media respectively, with >95% purity as determined by SDS-PAGE densitometry (Figure S1).

Actomyosin cosedimentation assays were performed to determine the fraction of M761-2R WT or S237C bound to phalloidin-stabilized F-actin under different metal conditions (Figure 1). Myosin and F-actin were brought to equilibrium before addition of metal-ATP and immediate centrifugation to capture the myosin fraction dissociated by ATP binding. In the absence of added nucleotide (apo), ≥ 95% of WT and S237C were bound to F-actin, consistent with a very low *K*_*d,actin*_ as previously reported (Kurzawa, Manstein, & Geeves, 1997) (Figure 1A, B). Upon addition of MgATP or MnATP, a large fraction of the WT protein detached from F-actin (*f*_*b*_ = 0.13 ± 0.04 and *f*_*b*_ = 0.20 ± 0.07, respectively), representing a weakening of actin binding by ∼100 fold (Figure 1B). In contrast, the S237C mutant retained relatively tight binding in the presence of MgATP (*f*_*b*_ = 0.68 ± 0.05), and detachment was rescued in the presence of MnATP (*f*_*b*_ = 0.22 ± 0.09). These results are consistent with previously published studies with M761 WT and S237C constructs without the α-actinin repeats, and confirm that the S237C substitution in M761-2R appears to modulate the actomyosin interaction base on the divalent metal present in solution (Cochran et al., 2013). This suggests that serine (-OH) in the WT switch-1 binds Mg^2+^ and Mn^2+^ sufficiently tightly to affect the release from F-actin, while the cysteine (-SH) in S237C can only bind Mn^2+^ well enough for efficient F-actin release.

**Figure 1.**
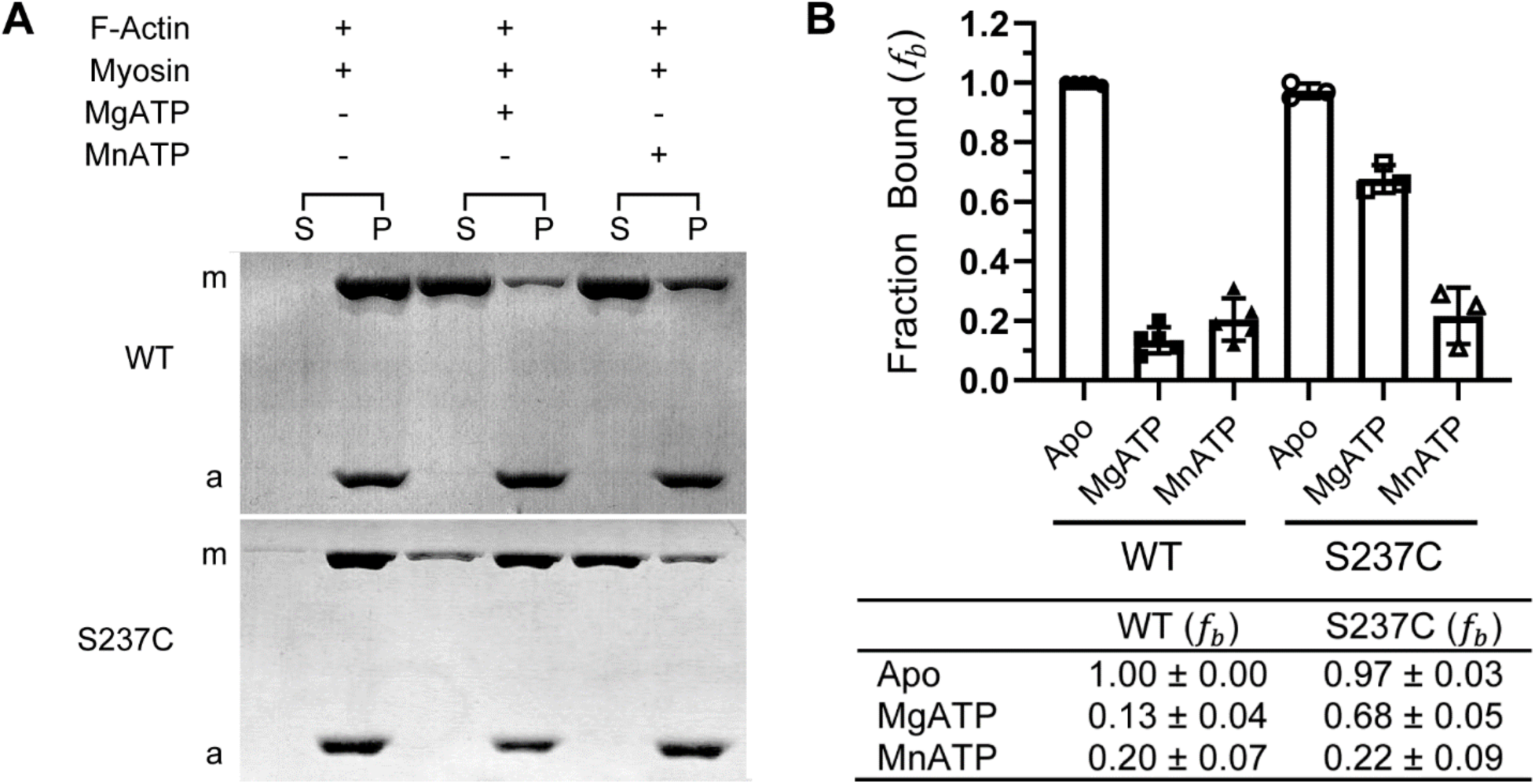
WT and S237C myosin cosedimentation with F-actin. **A**. Representative Coomassie Blue-stained SDS-PAGE gel of WT or S237C myosin (1 *µ*M) binding to phalloidin-stabilized F-actin (1.5 *µ*M). Equivalent volumes of the supernatant (S) and pellet (P) fractions were electrophoresed from each reaction and quantified by densitometry (B). Myosin is indicated with *m* and actin with *a*. **B**. Bar graph and table representing the mean fraction ± SD of WT (n=5) and S237C (n=3) myosin bound (*f*_*b*_) to F-actin.

### F-actin inhibition of the M761-2R S237C mutant with MnATP and CaATP is rescued by MnATP

To examine how changes in metal coordination affected catalytic activity, we performed NADH-coupled ATPase assays and determined the steady-state ATP turnover kinetics of WT and S237C M761-2R under different metal conditions (Figure 2). Our results with M761-2R constructs were consistent with previously published M761 data (Cochran et al., 2013). The ATPase rate of WT protein was slow in the presence of MgATP (0.02 ± 0.01 s^-1^) and showed weak activation in the presence of 10 *µ*M F-actin (0.08 ± 0.01 s^-1^) (Figure 2A and B). In contrast, WT ATP turnover was fast in the presence of Mn^2+^ (1.00 ± 0.13 s^-1^) and Ca^2+^ (0.31 ± 0.14 s^-1^), with no statistically significant F-actin stimulation, consistent with a weak F-actin affinity.

**Figure 2.**
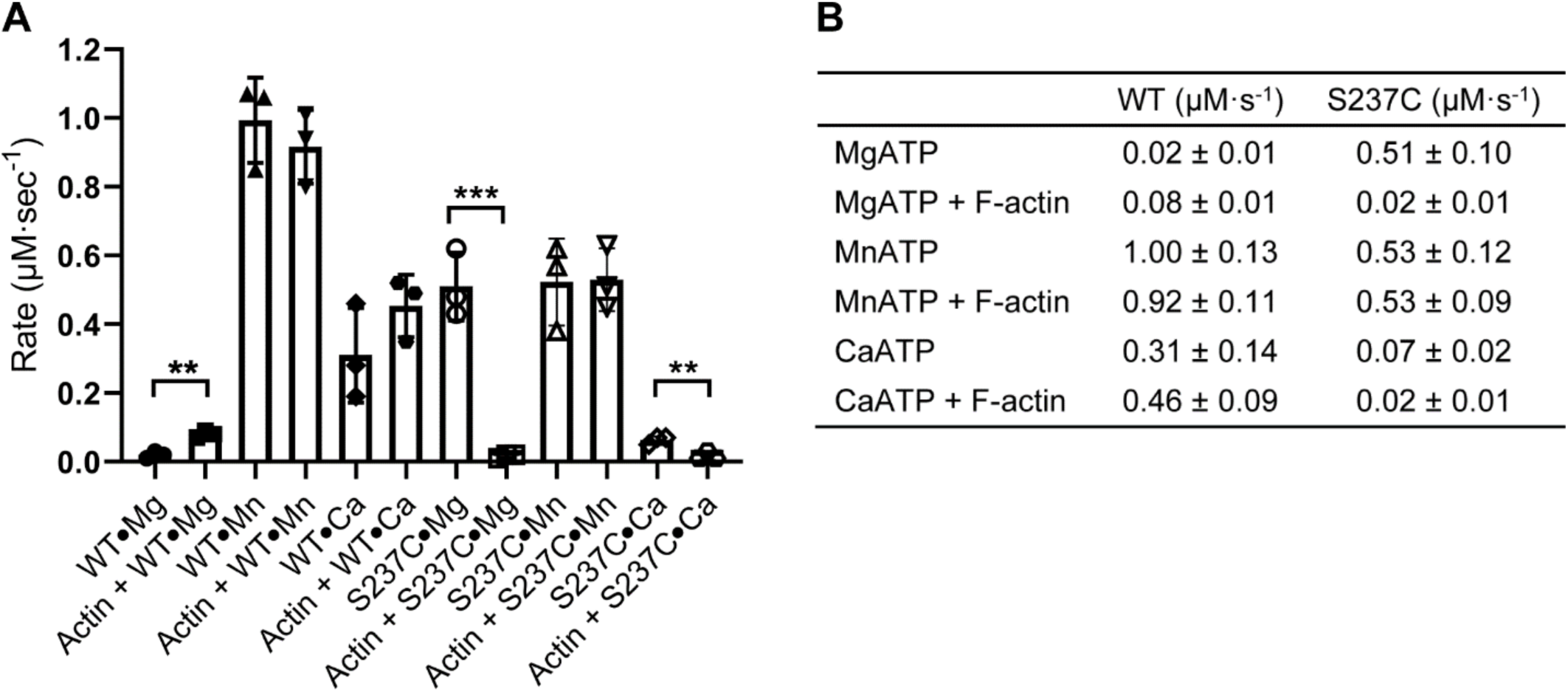
Steady-state ATPase activity of WT and S237C myosin. The NADH-coupled assay was used to determine the ATPase activity of 0.3 *µ*M WT or S237C myosin with 1 mM ATP and 1 mM divalent metal (Mg^2+^, Mn^2+^, or Ca^2+^) in the presence or absence of 20 *µ*M phalloidin-stabilized F-actin (n=3). **A**. Bar graph of the mean ATP turnover rates ± SD with different divalent metals. Unpaired *t*-tests were performed for each condition ± actin. **, p ≤ 0.01; ***, p ≤ 0.001. **B**. Table of the mean ± SD rate of ATP turnover for each myosin and condition.

Previous studies found that divalent metals have an inhibitory effect on myosin basal ATPase (Malik, Marchioli, & Martonosi, 1972), yet Mg^2+^ stimulates the actomyosin complex ATPase activity (Burke, Reisler, & Harrington, 1973; Kobayashi, Ramirez, & Warren, 2019). The inhibitory effect of divalent metals on basal ATPase rate is likely due to inhibition of product release, given that the metal is coordinated by both the hydrolysis products and the switch-loops (Swenson et al., 2014). Therefore, the fast basal ATPase rates with Mn^2+^ and Ca^2+^ that we report here and elsewhere (Cochran et al., 2013) are probably caused by a much more dynamic switch-1 loop at the active site due to weakened loop-divalent metal interactions.

In contrast to WT M761-2R, the S237C mutant ATPase was fast in the presence of MgATP (0.51 ± 0.10 s^-1^) but was strongly inhibited by F-actin (0.02 ± 0.01 s^-1^) (Figure 2A and B). In MnATP, the S237C mutant ATPase was also fast in the absence (0.53 ± 0.12 s^-1^) or presence of F-actin (0.53 ± 0.09 s^-1^), thus rescuing F-actin stimulated ATPase. On the other hand, the mutant CaATPase activity was slow (0.07 ± 0.02 s^-1^) and was inhibited by F-actin (0.02 ± 0.01 s^-1^).

The fast basal ATPase rates in the S237C mutant that we observed supports our hypothesis in that the mutant has a weak divalent metal coordination of switch-1 resulting in fast opening and closing of the nucleotide pocket causing high ATPase activity. F-actin inhibition of ATP hydrolysis in MgATP is likely explained by the switch loops being stabilized in an open conformation when F-actin is bound, and thereby inhibiting switch-1 closure in the very weak metal interactions (i.e. S237C with Mg^2+^). MnATP rescue of F-actin ATPase for the S237C mutant combined with the efficient F-actin release observed in our cosedimentation assays (Figure 1) indicate that in contrast to Mg^2+^, the switch-1 and Mn^2+^ divalent metal interaction is strong enough to close switch-1 when F-actin is bound and promote the conformational change at the F-actin binding site.

### F-actin gliding is divalent metal dependent in WT and abolished in S237C myosin

F-actin gliding assays were performed to determine the effect of the S237C mutation and metal conditions on force generation (Figure 3A). In the absence of divalent metal or ATP, no actin movement was observed (data not shown), consistent with the need for coupled ATPase activity and F-actin binding for motility. Robust F-actin motility was observed for WT M761-2R in the presence of MgATP (0.11 ± 0.01 *µ*m/s), in good agreement with previously published work (Anson et al., 1996) (Figure 3B and C). Interestingly, in the presence of MnATP the motility for WT was inhibited approximately three-fold (0.04 ± 0.01 *µ*m/s), and motility was abolished in the presence of CaATP (Figure 3B and C). The S237C mutant protein did not produce F-actin gliding under any condition (Figure 3B and C), and inhibited F-actin gliding when mixed with WT protein (Table S1).

**Figure 3.**
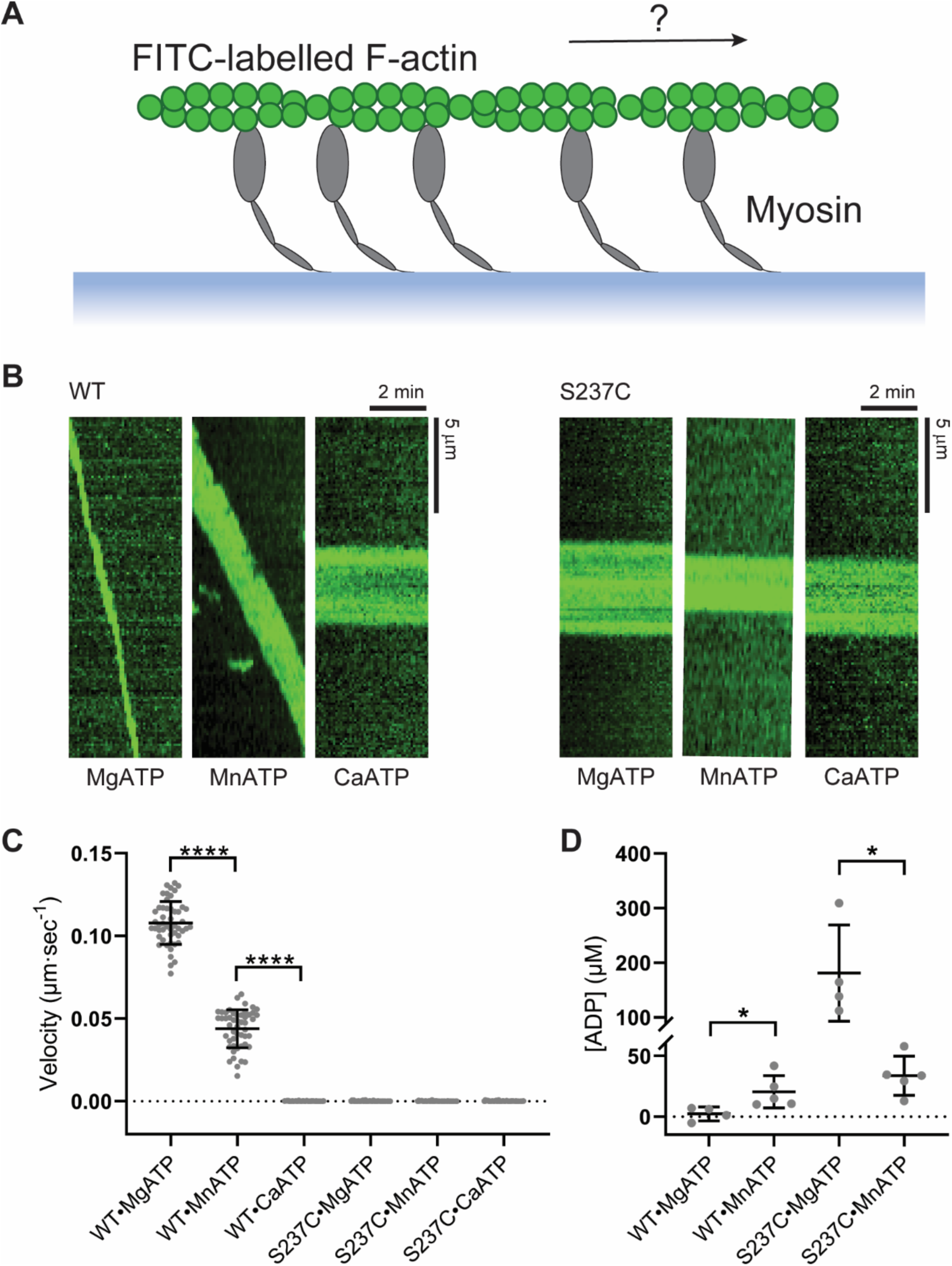
F-actin gliding by WT and S237C myosin under various metal conditions. **A**. Schematic showing myosin immobilized on a microscope glass chamber and FITC-labelled F-actin bound to myosin before addition of divalent metal and ATP to induce gliding. **B**. Representative kymographs of WT and S237C bound F-actin filaments in the different ATP and divalent metal compositions. **C**. Scatterplot of the gliding velocity of individual actin filaments with the mean ± SD indicated. For each condition, gliding velocities were calculated from the slope of 10 filament kymographs per gliding chamber from five separate experiments (n=50). WT(MgATP) = 0.11 ± 0.01 *µ*m·s^-1^, WT(MnATP) = 0.04 ± 0.01 *µ*m·s^-1^. For all other conditions gliding was not detected. An unpaired *t*-test was performed to test significance between the indicated conditions. ****, p ≤ 0.0001. **D**. F-actin gliding chambers were filled with the NADH-coupled system and incubated at 25°C for 45 min before the solution was extracted and the amount of ADP produced measured. The amount of ADP produced in each chamber is graphed as a scatter plot with the mean ± SD indicated. WT(MgATP) = 2 ± 6 *µ*M (n=4), WT(MnATP) = 20 ± 13 *µ*M (n=5), S237C(MgATP) = 181 ± 88 *µ*M (n=4), S237C(MnATP) = 34 ± 16 *µ*M (n=5). An unpaired *t*-test was performed to test significance between the indicated conditions. *, p < 0.05.

To ensure that the inhibition of motility was not simply due to inactivation of the enzyme in the chamber, we measured the ATPase activity of WT and S237C in the gliding chambers using the NADH-coupled assay by retrieving the chamber solution and measuring the ADP concentration (Figure 3D). As a control we first observed F-actin gliding with WT M761-2R in a subset of experiments before obtaining the chamber solution to measure the ADP concentration. Given the low fraction of filament bound myosin in the gliding chamber, we expect the overall ATPase rates to resemble basal activity. Consistent with the slow ATPase rates measured in solution-based assays (Figure 2), the WT enzyme had very little ADP build up (2 ± 6 *µ*M) in the presence of MgATP compared to MnATP (23 ± 14 *µ*M; p <0.05). The results for the S237C mutant were also consistent with the solution-based ATPase rates (Figure 2) in that in the presence of MgATP and MnATP large amounts of ADP were produced relative to the WT enzyme (181 ± 88 *µ*M and 34 ± 19 *µ*M; p <0.05) (Figure 3D). Our results show that both WT and S237C enzymes were enzymatically active in the gliding chambers, even in reduced or absent motility. In addition, this data supports the idea that WT is highly efficient (low ATPase, fast motility) in the presence of MgATP (Figure 4A) and shows a moderate degree of uncoupling of ATP hydrolysis to force generation in the presence of MnATP.

**Figure 4.**
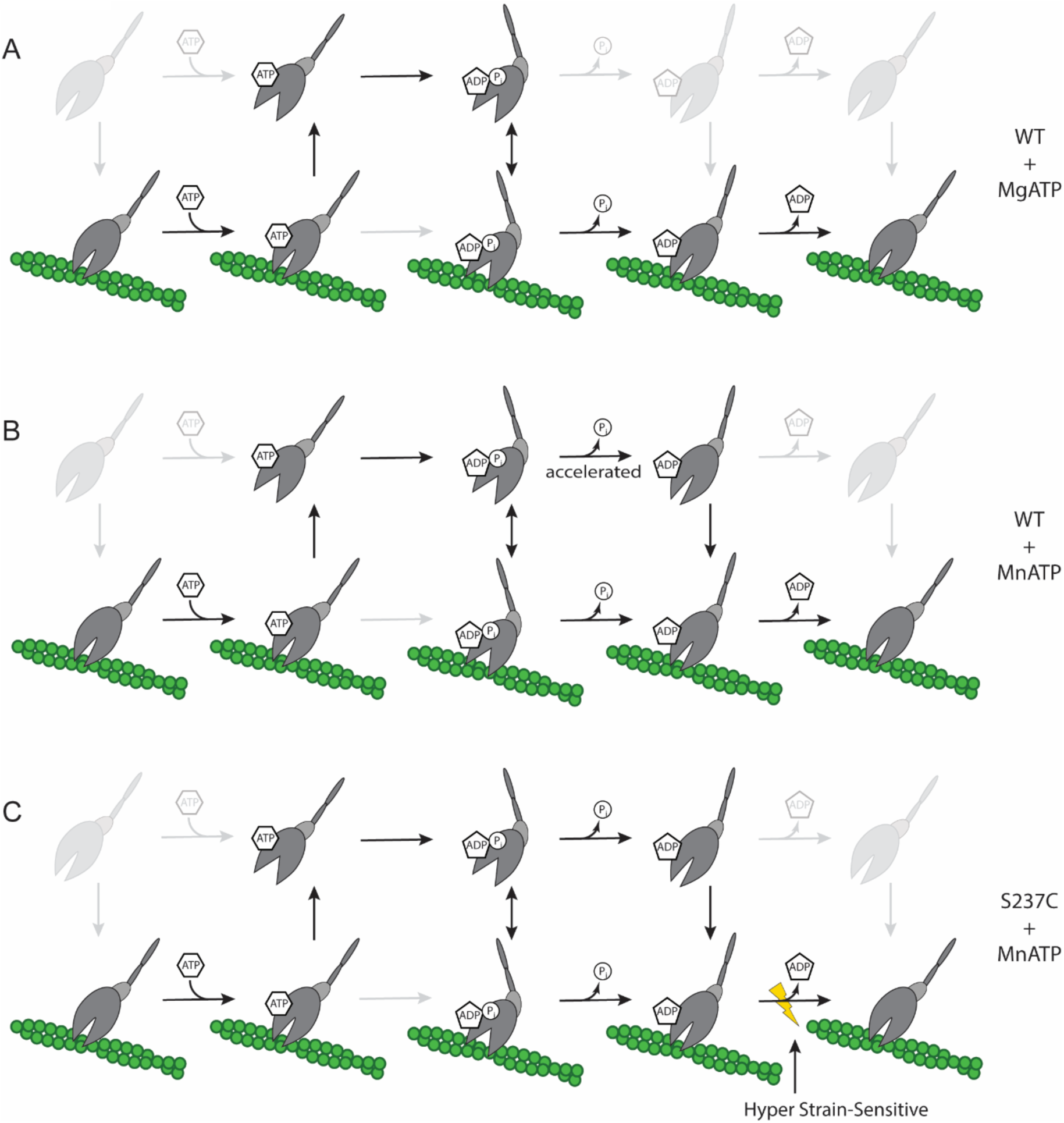
Models of ATPase cycle and force generation for myosin II. Kinetic models for myosin II motor constructs (grey) as they bind and hydrolyze ATP and release products in the presence of F-actin filaments (green). Main kinetic pathways are highlighted. **A**. Model for the WT MgATPase cycle. The conformational change that leads to tight F-actin binding intermediate, the power stroke, and P_i_ release limits both the basal and F-actin stimulated cycles (Gyimesi et al., 2008; Stein, Chock, & Eisenberg, 1984). Therefore, the dominant species in the ATPase cycle is the weak ADP-P_i_ state, and the power stroke occurs primarily when bound to F-actin (productive). **B**. Model for the WT MnATPase cycle. The conformational changes that are associated with P_i_ and ADP release limit the basal cycle while only the conformational change associated with P_i_ release limits the actomyosin cycle. Consequently, a significant fraction will undergo the power stroke off the filament (futile). **C**. Model for the S237C MnATPase cycle. The rate limiting steps are likely the same as WT (see panel B). ADP release is hyper strain sensitive and leads to inhibition of force generation along the F-actin filament.

The reduced motility of the WT construct and no motility of the S237C mutant construct in the presence of Mn^2+^ highlights the sensitivity of metal binding at the active site of myosin on force generation. It is already known that divalent metal binding significantly affects the ATPase cycle by changing the rate limiting step depending on the divalent metal at the active site (Cochran et al., 2013; Tkachev, Ge, Negrashov, & Nesmelov, 2013). Thus, even small perturbations of switch-1 loop allostery can significantly affect conformational changes throughout the motor, including the F-actin binding cleft and force generation elements. Switching metals directly destabilizes switch-1 coordination due to differences in affinity and atomic radii of the metal (Nihei & Tonomura, 1959) and may interfere with the salt bridge (R238:E459) between switch-1 and switch-2 loops, promoting faster phosphate release and increasing the rate of conformational changes in the motor (Tkachev et al., 2013). Premature loop movement and the associated effect on product release likely contribute to uncoupling of ATP turnover from force generation.

The mechanical elements linking switch loop movement and the power stroke (i.e. converter and lever arm) are unlikely interrupted in the presence of Mn^2+^ or Ca^2+^. Muscle fibers are able to contract and produce force in the presence of excess Ca^2+^ (albeit > 5-fold reduced), and rotational decay times showed differences between pre- and post-power stroke nucleotide states but no significant difference when these states were Ca^2+^ or Mg^2+^ coordinated (Polosukhina, Eden, Chinn, & Highsmith, 2000). Further evidence comes from crosslinking experiments where myosin S1 forms a 44 kDa converter to C-terminal crosslinked product only in the post-power stroke state but not in the pre-power stroke state (Pliszka & Karczewska, 2002). The most direct evidence comes from fluorescence studies where the bending or straightening of the converter in the pre- and post-power stroke could be measured using intrinsic fluorescence or FRET of added fluorophores. The transition from the post-power stroke to pre-power stroke can be directly observed in the presence of Mg^2+^, Mn^2+^, and Ca^2+^ (Ge, Gargey, Nesmelova, & Nesmelov, 2019; Tkachev et al., 2013). Consequently, the uncoupling effect of ATPase to force generation in the presence of Mn^2+^ or Ca^2+^ cannot readily be explained by a structural uncoupling in the motor domain.

While myosin likely undergoes lever arm movement during its MnATPase and CaATPase cycles, the rate of the conformational change associated with lever arm movement and phosphate release is significantly accelerated compared to the MgATPase cycle (Cochran et al., 2013; Ikebe, Inoue, & Tonomura, 1980; Sleep, Trybus, Johnson, & Taylor, 1981; Tkachev et al., 2013). Phosphate release is accelerated to the point that the conformational change associated with ADP release becomes rate limiting or co-rate limiting. Consequently, a significant proportion of myosin may undergo the power stroke and phosphate release before reattaching to the F-actin filament. The resulting increase in futile power strokes could explain the reduced motility observed for the WT in the presence of Mn^2+^ compared to Mg^2+^ (Figure 4A and B).

Interestingly, the S237C mutant was not motile (Figure 3) and surprisingly completely inhibited WT motility (Table S1), even in the presence of Mn^2+^. The rate limiting steps of the S237C basal MnATPase cycle were the same as WT (Cochran et al., 2013). In addition, the cosedimentation assays showed similar weak F-actin binding as the WT (Figure 1), suggesting a similar rate limiting step for the actomyosin ATPase cycle. Consequently, the lack of motility for S237C MnATPase cannot easily be explained by our solution-based assays alone. Unlike our solution-based ATPase and cosedimentation assays, myosin is under strain in the motility assays (Kron et al., 1991). Both muscle and non-muscle myosin II have strain sensitive ADP release steps where ADP release is slowed in the presence of strain, allowing prolonged tight F-actin binding to resist detachment (Kovacs, Thirumurugan, Knight, & Sellers, 2007; Mansson, 2010; Nyitrai & Geeves, 2004; Siththanandan, Donnelly, & Ferenczi, 2006; West et al., 2009). Therefore, at the high motor density the velocity of F-actin sliding in our motility assays was determined by the ADP release limited rate of actomyosin complex dissociation (Brizendine et al., 2017; Nyitrai et al., 2006; Siemankowski, Wiseman, & White, 1985; Weiss, Rossi, Pellegrino, Bottinelli, & Geeves, 2001; Yengo, Takagi, & Sellers, 2012). In addition, Mg^2+^ directly affects ADP release kinetics for myosin II, suggesting that divalent metal coordination by the switch loops may affect the strain sensitivity of ADP release (Chizhov, Hartmann, Hundt, & Tsiavaliaris, 2013; Swenson et al., 2014). We therefore propose that interfering with divalent metal coordination, by mutating switch-1 S237C, produced a more strain sensitive motor (Figure 4C). This exaggerated strain sensitivity may allow F-actin bound motors in the gliding assay to actively inhibit motility. Our model for the S237C mutant in the presence of Mn^2+^ is similar to WT in the presence of Mn^2+^, where a fraction of the motor undergoes futile power strokes (Figure 4C). However, the fraction of S237C undergoing the power stroke while bound to F-actin, and any other motors bound to the same filament, would experience exaggerated strain-dependent slowing of ADP release post-power stroke, which leads to an increase in motor-filament affinity and prolonged binding. This effect would be elevated when mixed with WT where strain is increased on the S237C motors when bound to the same filament. Therefore, we propose that exaggerated strain sensitivity of S237C may allow F-actin bound motors in the gliding assay to actively inhibit motility (Figure 4C). Future experiments will be key to testing our model and uncovering inhibition of motility by S237C.

F-actin dissociation (in solution) was significantly slowed for WT in the presence of Ca^2+^ and for S237C in the presence of Mg^2+^ and Ca^2+^ (Cochran et al., 2013; Ge et al., 2019; Tkachev et al., 2013). Under these conditions, there was unusually tight F-actin binding and F-actin inhibited ATP turnover (Figure 1, Figure 2, Cochran et al., 2013). We proposed previously that this inhibitory effect was due to the weakened interaction between switch-1 and divalent metal, slowing switch loop closure when the bound actin stabilizes the open conformation (Cochran et al., 2013). This explains dramatic weakening of ATP binding, which would cause a slow switch-1 induced opening of the F-actin cleft leading to the observed slowed ATP induced actomyosin dissociation. The slowing of F-actin dissociation would have an increased effect in motility assays where sliding velocity was already limited by the rate of detachment. Since detachment limits gliding velocity, it is unsurprising that higher duty ratios (time spent tightly bound to F-actin) correlate with slower motors (Bloemink & Geeves, 2011). We propose that the significantly slowed actin dissociation contributes to inhibition of motility in our assays, especially for WT in the presence of Ca^2+^.

It is worth noting that while the weakening of nucleotide binding and decrease in ATP-dependent actin dissociation was a general feature of WT CaATPase activity; the extent varies by construct and condition. The *D. discoideum* M761 construct has weakened nucleotide affinity to the extent that F-actin binding has a slight inhibitory effect (Cochran et al., 2013), while for the M758 construct, F-actin stimulates its CaATPase less than 2-fold (Korman, Anderson, Prochniewicz, Titus, & Thomas, 2006; Tkachev et al., 2013), and skeletal muscle myosin stimulation varies from as low as 1.5-fold to as high as 22-fold (Ge et al., 2019; Nihei & Tonomura, 1959; Peyser, Ajtai, Werber, Burghardt, & Muhlrad, 1997). In addition, muscle fibers are reported to produce (reduced) force in the presence of excess Ca^2+^ (Polosukhina et al., 2000). Consequently, other constructs likely vary in their duty ratios and possibly strain sensitivity in the presence of Ca^2+^ and may be capable of F-actin motility.

Together, our results strongly support the hypothesis that stable active site closure via switch-1 not only regulates ATP turnover and F-actin binding, but also regulates the coupling of ATP turnover to force generation along F-actin. Hence, interfering with metal coordination by exchanging divalent metals or mutation uncouples ATP hydrolysis from force generation by increasing the rate of ATP hydrolysis while inhibiting motility.

One interesting outcome of our study with myosin illustrates differences in the coupling between ATP hydrolysis and force production with the structurally related kinesin superfamily. Our observations show that while both myosins and kinesins share a similar fold and active site components (i.e. switch-1, switch-2, P-loop), their sensitivity to switch-1 divalent metal coordination and coupling to filament binding and productive force generation are clearly different. For kinesins, Mn^2+^ structurally and functionally replaced Mg^2+^ as a cofactor for ATP hydrolysis (Cochran, Zhao, Wilcox, & Kull, 2012). In WT kinesin, microtubule gliding velocities as well as basal and microtubule stimulated ATPase rates were similar with either Mn^2+^ or Mg^2+^, unlike in myosin where the basal ATPase activity was stimulated by MnATPbut not by MgATP. In addition, the switch-1 serine to cysteine mutant in kinesin is able to rescue ATP hydrolysis rates and partially rescue gliding velocity rates in the presence of Mn^2+^, in contrast to myosin where the cysteine mutant was unable to generate motility under any conditions. This suggests that filament binding and force generation in kinesins are less sensitive to switch-1 and divalent metal coordination than in myosins and that the coupling between these elements is less stable in myosins. Given the difference between the kinesin and myosin superfamilies, sensitivity of protein function to divalent metal coordination is likely specific among other P-loop containing NTPases. Since P-loop containing NTPases comprise the most abundant and diverse group of predicted gene products (Koonin, Wolf, & Aravind, 2000), probing the effect of divalent metal coordination on NTPase activity and biological functions using this metal-switch system is a promising approach to uncover mechanistic details of a large group of diverse proteins.

## MATERIALS AND METHODS

### Cloning, expression, and purification of M761(WT)-2R and M761(S237C)-2R

The pDXA-3H plasmid encoding the wild-type (WT) soluble head fragment (amino acids 2-761) of *Dictyostelium discoideum* myosin II with two α-actinin repeats, M761(WT)-2R (116.3 kDa), was a generous gift from the Manstein lab (Kliche, Fujita-Becker, Kollmar, Manstein, & Kull, 2001). To generate the myosin II soluble head fragment with the S237C mutation, M761(S237C) (Cochran et al., 2013) was digested with *Bam*HI and *Bst*XI, and the resulting fragment was ligated into the M761(WT)-2R background to generate M761(S237C)-2R. All clones were confirmed by sequencing. For protein expression, M761(WT)-2R and M761(S237C)-2R were transfected into *D. discoideum* AX3-ORF+ cells (dictyBase, Cat# DBS0235546, RRID: SCR_006643) using electroporation as described (Gaudet, Pilcher, Fey, & Chisholm, 2007). Subconfluent plates of transformed *D. discoideum* were used to seed 0.5 L of AX medium (Formedium, Cat# AXM0102) in 2 L flasks at 10^5^ cells/mL. Cultures were grown at 21°C and 120 rpm to 10^7^ cells/mL before harvesting by centrifugation.

The M761(WT)-2R was purified as described with modifications (Cochran et al., 2013). Specifically, cells from ∼1 L culture were resuspended in 60 mL of lysis buffer (50 mM Tris-HCl pH 8.0, 2 mM EDTA, 0.2 mM EGTA, 1 mM DTT, 5 mM benzamidine, 0.1 mM phenylmethylsulphonyl fluoride (PMSF), 1 tablet of cOmplete™ Protease Inhibitor Cocktail (Roche Applied Science, Cat# 11697498001), 2% (v/v) Triton X-100, 30 μg/ml RNase A, 1.7 U/mL units of alkaline phosphatase) and lysed for 15 min at 4°C. The cell lysate was clarified by centrifugation at 186,000 x *g* and 4°C for 45 min. The cytoskeletal pellet was homogenized with 100 mL of extraction buffer (50 mM Na-HEPES pH 7.3, 100 mM NaCl, 30 mM potassium acetate, 10 mM magnesium acetate, 5 mM benzamidine, 0.1 mM PMSF, 1 tablet of cOmplete™ Protease Inhibitor Cocktail). Clarification and homogenization were repeated with 20 mL extraction buffer containing 20 mM ATP to release the myosin motor from actin. A final 370,000 x *g* centrifugation at 4°C for 45 min yielded a myosin-containing supernatant that was loaded onto a 1 mL nickel-nitrilotriacetic acid agarose (Ni-NTA) (Qiagen, Cat# 30210) column equilibrated with column buffer (50 mM Tris-HCl pH 8.0, 150 mM NaCl, and 1 mM EDTA, 30 mM imidazole). The protein was eluted from the column with elution buffer (column buffer, 0.47 M imidazole) and adjusted to 5 mM EDTA to chelate trace divalent cations. The eluted protein was incubated at 4°C for 15 min before buffer exchange with a HiPrep 26/10 resin column into final buffer (20 mM Na-HEPES pH 7.3, 95 mM potassium acetate, 0.1 mM EDTA, 150 mM sucrose, and 1 mM DTT) and concentrated using a 10 kDa Amicon® centrifuge filter (Millipore, Cat# C7715). Purified protein was flash frozen in liquid nitrogen and stored at −80°C. For the M761(S237C) protein, 10 mM MnCl_2_ was added to the extraction buffer. Protein concentrations were determined using the Bradford Reagent (Sigma, Cat# B6916), SDS-PAGE densitometry, and absorption at A_280_ using 96,150 M^-1^cm^-1^ as the extinction coefficient.

### Preparation of F-actin

Chicken skeletal muscle G-actin was prepared from acetone powder as described (Spudich & Watt, 1971). G-actin was polymerized (F-actin) as described previously and stabilized by addition of phalloidin or fluorescein isothiocyanate (FITC) labelled phalloidin (Sigma Aldrich, Cat# P5282) in DMSO to a final molar concentration equal to or greater than F-actin (Cochran et al., 2013). Stabilized F-actin was pelleted at 214,000 x *g* for 40 min and resuspended in ATPase Buffer (20 mM Na-HEPES pH 7.2, 0.25 M KCl, 150 mM sucrose, 1 mM DTT).

### Actomyosin Cosedimentation Assays

Actomyosin cosedimentation experiments were performed by mixing myosin with phalloidin stabilized F-actin at the indicated concentrations in reaction buffer (20 mM Na-HEPES pH 7.2, 5 mM MgCl_2_, 250 mM KCl, 150 mM sucrose, 1 mM DTT) followed by incubation at 25°C for 10 min before the addition of either 10 mM MgATP or MnATP. After addition of nucleotide, solutions were immediately pelleted by centrifugation at 100,000 x *g* for 15 min at 25°C. Supernatants were removed, and pellets were resuspended in an equal volume of reaction buffer. Gel samples were prepared from equivalent volumes of supernatant and pellet fractions for each reaction by the addition of 5X Laemmli buffer (60 mM Tris-HCl pH 6.8, 0.4% w/v SDS, 10% v/v glycerol, 5% v/v β-mercaptoethanol, 0.01% w/v bromophenol blue) and analyzed by SDS-PAGE. Gels were stained with Coomassie, and densitometry was performed using the software FIJI (https://fiji.sc, RRID:SCR_002285) (Schindelin et al., 2012).

### Steady-state NADH-coupled ATPase assays

Steady-state ATPase activities were monitored using the NADH coupled assay in modified reaction buffer (20 mM Na-HEPES pH 7.2, 100 mM KCl, 150 mM sucrose, 1 mM DTT) at 25°C (Cruz, Sweeney, & Ostap, 2000; Hass, Boyer, & Reynard, 1961; Imamura, Tada, & Tonomura, 1966). Briefly, 0.2 *µ*M myosin in modified reaction buffer was mixed with an equal volume of NADH cocktail (1 mM ATP, 5 mM MgCl_2_ or MnCl_2_ or CaCl_2_, 0.4 mM NADH (Acros Organic, Cat# 271100010), 0.5 mM phosphoenolpyruvate (Tokyo Chemical Industry), 5 U/mL rabbit pyruvate kinase (Roche Diagnostics, Cat# 10128155001), 8 U/mL lactate dehydrogenase (Sigma-Aldrich)), and an equal volume of F-actin or modified reaction buffer. After mixing, reactions were transferred into a 384-well plate in duplicate, pulse centrifuged at 1000 x *g*, and placed in a microplate spectrophotometer (BioTek) where NADH oxidation was monitored at 340 nm over time. The amount of NADH oxidized (which equals the amount of ADP produced) was calculated using an NADH standard curve (0 *µ*M – 800 *µ*M NADH). The NADH stocks and standard curves were confirmed by using the extinction coefficient of NADH (6220 M^-1^cm^-1^) at 340 nm.

### F-actin gliding assays

Gliding assays were performed as described with modifications (Kron, Toyoshima, Uyeda, & Spudich, 1991). All experiments were performed in motility buffer, which contained either 20 mM MOPS pH 7.5 or 25 mM Na-HEPES pH 7.5, 25 mM KCl, 1 mM EGTA, 1 mM DTT. HEPES was used instead of MOPS after determining it had no effect on motility (data not shown). Incubations were performed in a humid container at 4°C. Briefly, myosin (50 – 100 *µ*g/mL) was infused in a nitrocellulose treated glass chamber and incubated for 5 min. Flow chambers held ∼10 μL of solution. An additional two chamber volumes of myosin were wicked through the chamber and incubated for ≥ 5 min. The flow chamber was washed with two chamber volumes of blocking buffer (motility buffer, 0.5 mg/mL BSA) and incubated for 1 min. Two chamber volumes of F-actin (motility buffer, 0.5 mg/mL BSA, 0.01 *µ*M FITC-labelled F-actin) were then wicked through the chamber and incubated for 2 min. Excess F-actin was washed out of the chamber by infusing two chamber volumes of wash buffer (motility buffer, 0.5 mg/mL BSA, oxygen scavenger system (25 mM glucose, 0.2 mg/mL glucose oxidase, 175 *µ*g/mL catalase, 71.6 mM β-mercaptoethanol) at 25°C. Static F-actin filaments were imaged using a Nikon NiE epifluorescence microscope equipped with a Plan Apo VC 60X 1.4 NA oil objective and a Hamamatsu Orca-Flash 2.8 sCMOS camera controlled by Nikon Elements software. To activate gliding, chambers were infused with two chamber volumes of gliding buffer (motility buffer, oxygen scavenger system, 20 U/mL pyruvate kinase, 2.5 mM phosphoenolpyruvate, 0.3 % w/v methylcellulose (4,000 cP), 2 mM ATP, 2 mM of divalent metal (MgCl_2_, MnCl_2_, or CaCl_2_)) and sealed with VALAP. Time-lapse images were recorded in 2-10 sec intervals. Images and movies were acquired with identical exposures and scaled identically in FIJI.

## ACKNOWLEDGEMENTS

We thank the Manstein Lab for their generous provision of the pDXA-3H:M761-2R plasmid. We thank Stephanie Ems-McClung and members of the Walczak Lab for valuable discussion, comments, and guidance on the manuscript. We thank the dictyBase for providing the *Dictyostelium discoideum* cell line as well as relevant references for protocols. We thank the Yu-Li Wang lab for making their actin purification protocols available to us. This work is supported by NSF Grant MCB 1614514 to JCC and CEW. Some of the microscopy was carried out in the IU Light Microscopy Imaging Center, which is supported in part by the Office of the Vice Provost for Research, College of Arts and Sciences, School of Medicine, Indiana University, Bloomington, and the School of Optometry at Indiana University.

