## Supplemental Figure and Table for "Switch-1 Instability at the Active Site Decouples ATP Hydrolysis from Force Generation in Myosin II"

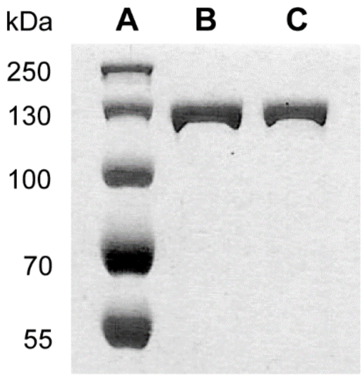


**Figure S1. Coomassie-Blue stained SDS-PAGE gel of purified WT and S237C myosin M761-2R proteins.** Lane A, PageRuler Plus protein ladder. Lanes B and C, 31 pmol of WT (B) and S237C (C) proteins.

| WT / S237C | Condition | Velocity (μm/sec) |
|---|---|---|
| + / – | MgATP | 0.11 ± 0.02 |
| + / + | MgATP | no motility |
| + / – | MnATP | 0.04 ± 0.01 |
| + / + | MnATP | no motility |

**Table S1. F-actin gliding with mixtures of WT and S237C myosin.** F-actin gliding velocity of individual actin filaments with the mean ± SD indicated of WT with and without S237C under MgATP (n=2) and MnATP (n=3) conditions. A minimum of 10 F-actin filaments were tracked per condition per experiment. Experiments were performed as described with WT and S237C mixtures varying from 1:1 to 5:2 ratios.
